# P38α Regulates Expression of DUX4 in Facioscapulohumeral Muscular Dystrophy

**DOI:** 10.1101/700195

**Authors:** L. Alejandro Rojas, Erin Valentine, Anthony Accorsi, Joseph Maglio, Ning Shen, Alan Robertson, Steven Kazmirski, Peter Rahl, Rabi Tawil, Diego Cadavid, Lorin A. Thompson, Lucienne Ronco, Aaron N. Chang, Angela M. Cacace, Owen Wallace

## Abstract

FSHD is caused by the loss of repression at the *D4Z4* locus leading to DUX4 expression in skeletal muscle, activation of its early embryonic transcriptional program and muscle fiber death. While progress toward understanding the signals driving DUX4 expression has been made, the factors and pathways involved in the transcriptional activation of this gene remain largely unknown. Here, we describe the identification and characterization of p38α as a novel regulator of DUX4 expression in FSHD myotubes. By using multiple highly characterized, potent and specific inhibitors of p38α/β, we show a robust reduction of DUX4 expression, activity and cell death across FSHD1 and FSHD2 patient-derived lines. RNA-seq profiling reveals that a small number of genes are differentially expressed upon p38α/β inhibition, the vast majority of which are DUX4 target genes. Our results reveal a novel and apparently critical role for p38α in the aberrant activation of DUX4 in FSHD and support the potential of p38α/β inhibitors as effective therapeutics to treat FSHD at its root cause.

**VISUAL ABSTRACT:** 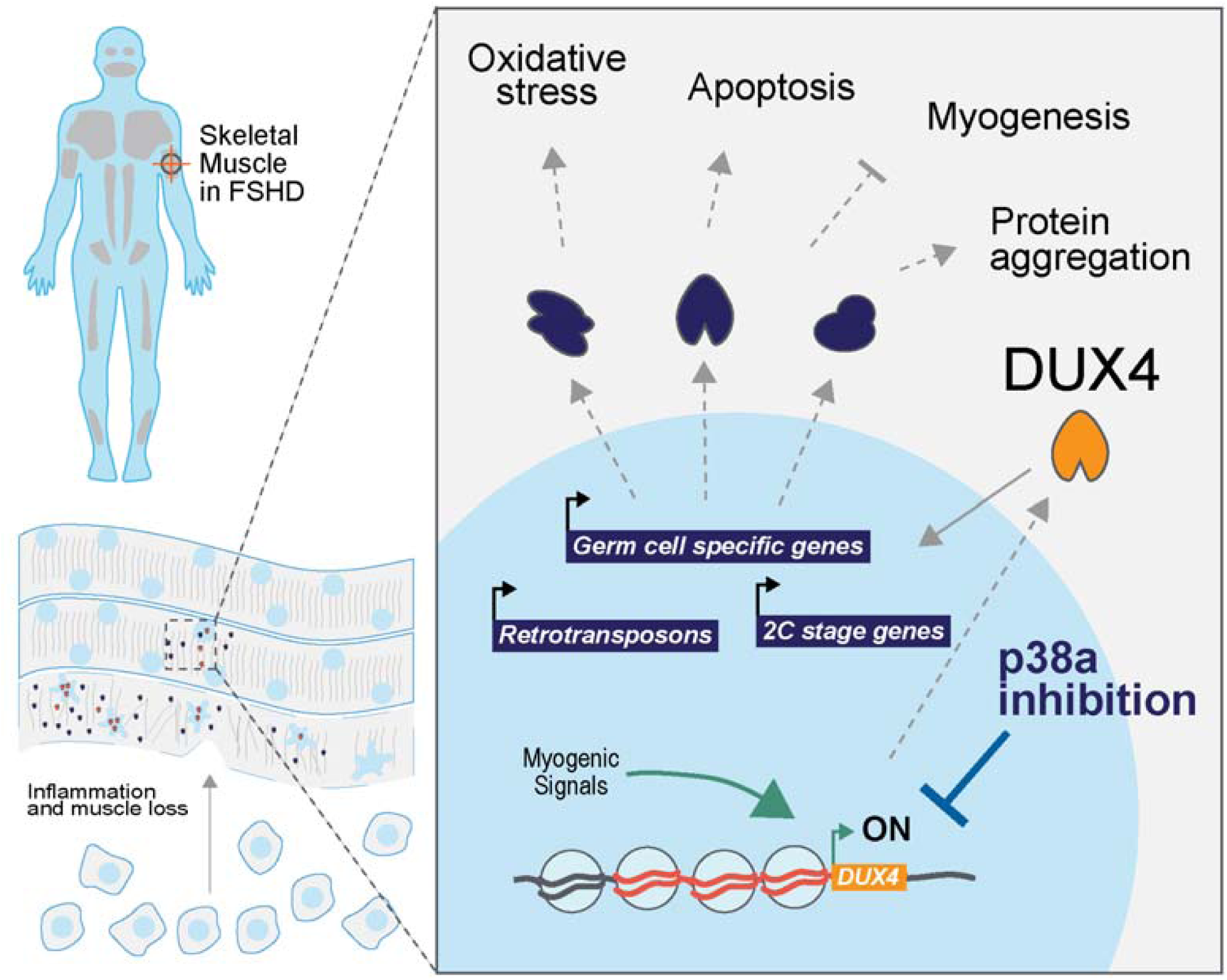

## INTRODUCTION

Facioscapulohumeral muscular dystrophy (FSHD) is a rare and disabling condition with an estimated worldwide population prevalence of between 1 in 8,000-20,000 (Statland and Tawil, 2014; Deenen *et al*., 2014). Most cases are familial and inherited in an autosomal dominant fashion and about 30% of cases are known to be sporadic. FSHD is characterized by progressive skeletal muscle weakness affecting the face, shoulders, arms, and trunk, followed by weakness of the distal lower extremities and pelvic girdle. Initial symptoms typically appear in the second decade of life but can occur at any age resulting in significant physical disability in later decades (Tawil *et al*., 2015). There are currently no approved treatments for this condition.

FSHD is caused by aberrant expression of the *DUX4* gene, a homeobox transcription factor in the skeletal muscle of patients. This gene is located within the *D4Z4* macrosatellite repeats on chromosome *4q35*. DUX4 is not expressed in adult skeletal muscle when the number of repeat units (RU) is >10 and the locus is properly silenced (Lemmers *et al*., 2010). In most patients with FSHD (FSHD1), the *D4Z4* array is contracted to 1–9 RU in one allele. FSHD1 patients carrying a short a *D4Z4* (1–3 RU) are on average more severely affected than those with longer array (8-9) (Tawil *et al*., 1996). Loss of these repetitive elements leads to de-repression of the *D4Z4* locus and ensuing aberrant *DUX4* expression in skeletal muscle (de Greef *et al*., 2009; Wang *et al*., 2018). In FSHD2, patients manifest similar signs and symptoms as described above but genetically differ from FSHD1. These patients have longer *D4Z4* arrays but exhibit similar de-repression of the locus with low levels of DNA methylation (Jones *et al*., 2014; 2015; Calandra *et al*., 2016). This loss of repression is caused by mutations in *SMCHD1,* an important factor in the proper deposition of DNA methylation across the genome (Jansz *et al*., 2017; Dion *et al*., 2019). *SMCHD1* has also been identified as the cause of Bosma arhinia microphthalmia syndrome (BAMS), a rare condition characterized by the lack of an external nose (Shaw *et al*., 2017; Gordon *et al*., 2017; Mul *et al*., 2018). Similarly, modifiers of the disease, such as *DNMT3B*, are thought to participate in the establishment of silencing (van den Boogaard *et al*., 2016).

*DUX4* expression in skeletal muscle as a result of the *D4Z4* repeat contraction or *SMCHD1* mutations leads to activation of a downstream transcriptional program that causes FSHD (Yao *et al*., 2014; Bosnakovski *et al*., 2014; Homma *et al*., 2015; Jagannathan *et al*., 2016; Shadle *et al*., 2017). Major target genes of DUX4 are members of the DUX family itself and other homeobox transcription factors. Additional target genes include highly homologous gene families, including the preferentially expressed in melanoma (PRAMEF), tripartite motif-containing (TRIM) and methyl-CpG binding protein-like (MBDL) (Geng *et al*., 2011; Tawil *et al*., 2014; Yao *et al*., 2014; Shadle *et al*., 2017). Expression of DUX4 and its downstream transcriptional program in skeletal muscle cells is toxic, leading to dysregulation of multiple pathways resulting in impairment of contractile function and cell death (Bosnakovski *et al*., 2014; Tawil *et al*., 2014; Homma *et al*., 2015; Rickard *et al*., 2015; Himeda *et al*., 2015; Statland *et al*., 2015).

Several groups have made progress towards understanding the molecular mechanisms regulating DUX4 expression (van den Boogaard *et al*., 2015; van den Boogaard *et al*., 2016; Campbell *et al*., 2018; Oliva *et al*., 2019). However, factors that drive transcriptional activation of DUX4 in FSHD patients are still largely unknown. By screening our annotated chemical probe library to identify disease-modifying small molecule drug targets that reduce DUX4 expression in FSHD myotubes, we have identified multiple chemical scaffolds that inhibit p38 α and β mitogen-activated protein kinase (MAPK). We found that inhibitors of p38α kinase or its genetic knockdown, reduce DUX4 and its downstream gene expression program in FSHD myotubes, thereby impacting the core pathophysiology of FSHD.

Members of the p38 MAPK family, composed of α, β, γ and δ, isoforms are encoded on separate genes and play a critical role in cellular responses needed for adaptation to stress and survival (Whitmarsh, 2010; Krementsov *et al*., 2013; Martin *et al*., 2015). In many inflammatory, cardiovascular and chronic disease states, p38 MAPK stress-induced signals can trigger maladaptive responses that aggravate, rather than alleviate, the disease process (Martin et al., 2015; Whitmarsh, 2010). Similarly, in skeletal muscle, a variety of cellular stresses including chronic exercise, insulin exposure and altered endocrine states, myoblast differentiation, reactive oxygen species as well as apoptosis have all been shown to induce the p38 kinase pathways (Zarubin and Han, 2005; Keren *et al*., 2006). Moreover, these pathways can be activated by several external stimuli, including pro-inflammatory cytokines and cellular stress environments, that lead to activation of the upstream kinases MKK3 and MKK6. Activation of these, which in turn phosphorylate p38 in its activation loop, trigger downstream phosphorylation events. These include phosphorylation of other kinases, downstream effectors like HSP27 and transcription factors culminating in gene expression changes (Kyriakis and Avruch, 2001; Viemann *et al*., 2004; Cuenda and Rousseau, 2007).

P38α is the most abundantly expressed isoform in skeletal muscle and it has an important role controlling the activity of transcription factors that drive myogenesis (Simone *et al*., 2004; Knight *et al*., 2012; Segalés *et al*., 2016). P38α abrogation in mouse myoblasts inhibits fusion and myotube formation in vitro (Zetser *et al*., 1999; Perdiguero *et al*., 2007). However, conditional ablation of p38α in the adult mouse skeletal muscle tissue appears to be well-tolerated and alleviates phenotypes observed in models of other muscular dystrophies (Wissing *et al*., 2014).

Here, we show that selective p38α/β inhibitors potently decrease the expression of DUX4, its downstream gene program and cell death in FSHD myotubes across a variety of FSHD1 and FSHD2 genotypes. Using RNA-seq and high content image analysis we also demonstrated that myogenesis is not affected at concentrations that result in downregulation of DUX4.

## MATERIALS AND METHODS

### Cell lines and cell culture

Immortalized myoblasts from FSHD (AB1080FSHD26 C6) and healthy individuals (AB1167C20FL) were generated and obtained from the Institut Myologie, France. In short, primary myoblast cultures were obtained from patient samples and immortalized by overexpression of TERT and CDK4 (Krom *et al*., 2012). Primary myoblasts were isolated from FSHD muscle biopsies and were obtained from University of Rochester.

Immortalized myoblasts were expanded on gelatin-coated dishes (EMD Millipore, #ES-006-B) using Skeletal muscle cell growth media (Promocell, #C-23060) supplemented with 15% FBS (ThermoFisher, #16000044). Primary myoblasts were also expanded on gelatin-coated plates but using media containing Ham’s F10 Nutrient Mix (ThermoFisher, #11550043), 20% FBS and 0.5% Chicken embryo extract (Gemini Bio-product, #100-163P). For differentiation, immortalized or primary myoblasts were grown to confluency in matrigel-coated plates (Corning, #356234) and growth media was exchanged for differentiation media (Brainbits, #Nb4-500) after a PBS wash. DMSO (vehicle) or compounds (previously dissolved in DMSO at 10 mM stock concentrations) were added at the desired concentration at the time differentiation media was exchanged and maintained in the plates until harvesting or analysis.

### Small molecule compounds and antisense oligonucleotides

SB239063, Pamapimod, LY2228820 and Losmapimod were purchased from Selleck Chem (#S7741, S8125, S1494 and S7215). 10 mM stock solutions in DMSO were maintained at room temperature away from light. DUX4 antisense oligonucleotides (gapmer) were purchased from QIAGEN and were designed to target exon 3 of DUX4. The lyophilized oligos were resuspended in PBS at 25 mM final concentration and kept frozen at -20°C until used. This antisense oligonucleotide was added to cells in growth media 2 days before differentiation and maintained during the differentiation process until harvesting.

### Detection of DUX4 and target gene expression by RT-qPCR

RNA from myotubes was isolated from C6 FSHD cells differentiated in 6-well plates using 400 μl of tri-reagent and transfer to Qiagen qiashredder column (cat#79656). An equal amount of 100% Ethanol was added to flow through and transferred to a Direct-zol micro column (Zymo research cat# 2061) and the manufacturers protocol including on-column DNA digestion was followed. RNA (1 μg) was converted to cDNA using Superscript IV priming with oligo-dT (Thermofisher cat# 18091050). Pre-amplication of DUX4 and housekeeping gene *HMBS* was performed using preamp master mix (Thermofisher cat#4384267) as well as 0.2X diluted taqman assays (IDT DUX4 custom; forward Forward: 5’-GCCGGCCCAGGTACCA-3’, Reverse: 5’-CAGCGAGCTCCCTTGCA-3’, and Probe: 5’-/56-FAM/CAGTGCGCA/ZEN/CCCCG/3IABkFQ/-3’; and *HMBS* HS00609297m1-VIC). After 10 cycles of pre-amplification, reactions were diluted 5-fold in nuclease-free water and qPCR was performed using taqman multiplex master mix (Thermofisher cat#4461882).

To measure DUX4 target gene expression in a 96-well plate format, cells were lysed into 25 μL Realtime Ready lysis buffer (Roche, #07248431001) containing 1% RNAse inhibitor (Roche, #03335399001) and 1% DNAse I (ThermoFisher, #AM2222) for 10 min while shaking on a vibration platform shaker (Titramax 1000) at 1200 rpm. After homogenization, lysates were frozen at -80°C for at least 30 min and thawed on ice. Lysates were diluted to 100 μL using RNase-free water. 1 μL of this reaction was used for reverse transcription and preamplification of cDNA in a 5 μL one-step reaction using the RT enzyme from Taqman RNA-to-Ct (ThermoFisher, #4392938) and the Taqman Preamp Master Mix (ThermoFisher, #4391128) according to manufacturer’s specifications. This preamplification reaction was diluted 1:4 using nuclease-free water, 1μL of this reaction was used as input for a 5 μL qPCR reaction using the Taqman Multiplex Master Mix (ThermoFisher, #4484262). Amplification was detected in a Quantstudio 7 Flex instrument from ThermoFisher. The following Taqman probes were purchased from ThermoFisher; MBD3L2 Taqman Assay (ThermoFisher, Hs00544743_m1, FAM-MGB). ZSCAN4 Taqman Assay (ThermoFisher, Hs00537549_m1, FAM-MGB). LEUTX Taqman Assay (Thermo Fisher, Hs01028718_m1, FAM-MGB). TRIM43 Taqman Assay (ThermoFisher, Hs00299174_m1, FAM-MGB). KHDC1L Taqman Assay (ThermoFisher, Hs01024323_g1, FAM-MGB). POLR2A Taqman Assay (ThermoFisher, Hs00172187_m1, VIC-MGB).

### Detection of HSP27 by Electrochemiluminescence

Total and phosphorylated HSP27 was measured using a commercial MesoScale Discovery assay, Phospho (Ser82)/Total HSP27 Whole Cell Lysate Kit (MesoScale Discovery, # K15144D). Myotubes were grown in 96-well plates using conditions described above and were lysed using 25 μL of 1X MSD lysis buffer with protease and phosphatase inhibitors. The lysates were incubated at room temp for 10 minutes with shaking at 1200 rpm using Titramax 1000. Lysates were stored at -80 °C until all timepoints were collected. Lysates were then thawed on ice and 2 μL were used to perform a BCA protein assay (ThermoFisher, # 23225). 10 μL of lysate were diluted 1:1 in 1X MSD lysis buffer and added to the 96-well Mesoscale assay plate. Manufacturer instructions were followed, and data was obtained using a MesoScale Discovery SECTOR S 600 instrument.

### Myotube nuclei isolation and detection of DUX4 by Electrochemiluminescence

DUX4 was measured using a novel MesoScale Discovery assay developed at Fulcrum Therapeutics. Anti-DUX4 monoclonal capture antibody (clone P2B1) was coated overnight at 5 μg/ml in 0.1 M sodium Bicarbonate pH=8.4 onto a Mesoscale 384 well plate (L21XA). The plate was blocked with 5% BSA/PBS for at least 2 hours. Human FSHD myotubes grown in 100 mm plates in the conditions described above were harvested 4 days post differentiation using TrypLE express solution (Gibco, #12605-010), neutralized with growth media and the myotubes were pelleted by centrifugation. Myotubes were resuspended in ice cold nuclei extraction buffer (320 mM Sucrose, 5 mM MgCl2, 10 mM HEPES, 1% Triton X-100 at pH=7.4). Nuclei were pelleted by centrifugation at 2000 xg for 4 minutes at 4°C. Nuclei were resuspended in ice cold wash buffer (320 mM Sucrose, 5 mM MgCl2, 10 mM HEPES at pH=7.4) and pelleted by centrifugation at 2000 xg for 4 minutes at 4°C. Nuclei were suspended in 150 μl of RIPA buffer at 4°C (+150 mM NaCl). Extracts were diluted 1:1 with assay buffer and 10 μl per well was added to 384 well pre-coated/blocked MSD plate and incubated for 2 hours. Anti-DUX4-Sulfo Conjugate (clone E5-5) was added to each well and incubated for two hours. Plates were washed and 40 μl per well of 1X Read T buffer was added. Data was obtained using a MesoScale Discovery SECTOR S 600 instrument.

### Quantitative Immunofluorescent detection of Myosin Heavy Chain, SLC34A2 and cleaved Caspase-3

Myotubes were grown and treated as described above. At day 5 after differentiation was induced, cells were fixed using 4% paraformaldehyde in PBS during 10 min at room temperature. Fixative was washed, and cells were permeabilized using 0.5% Triton X-100 during 10 min at room temperature. After washing, fixing and permeabilizing, the cells were blocked using 5% donkey serum in PBS/0.05% Tween 20 during 1 h at room temperature. Primary antibodies against MHC (MF20, R&D systems, #MAB4470), SLC34A2 (Cell signaling, #66445) and active Caspase-3 (Cell signaling, #9661) were diluted 1:500 in PBS containing 0.1% Triton X-100 and 5% donkey serum and incubated with cells for 1 h at room temperature. After 4 washes, secondary antibodies were added (ThermoFisher, #A32723 and # R37117) in a 1:2000 dilution and incubated during 1 h at room temperature. During the last 5 min of incubation a 1:2000 dilution of DAPI was added before proceeding with final washes and imaging. Images were collected using the CellInsight CX7 (ThermoFisher). Images were quantified using HCS Studio Software. Differentiation was quantified by counting the percentage of nuclei in cells expressing MHC from the total of the well. SLC34A2 and active Caspase-3 signal was quantified by colocalization of cytoplasmic cleaved Caspase-3 within MHC expressing cells.

### Knockdown of *MAPK12* and *MAPK14* in FSHD myotubes

Exponentially dividing immortalized C6 FSHD myoblasts were harvested and counted. 50000 myoblasts were electroporated using a 10 μL tip in a Neon electroporation system (ThermoFisher). Conditions used were determined to preserve viability and achieved maximal electroporation (Pulse V=1100V, pulse width=40 and pulse #=1). After electroporation, cells were plated in growth media and media was changed for differentiation 24h after. 3 days after differentiation, cells were harvested and analyzed for KD and effects in *MBD3L2* using the RT-qPCR assay described before. siRNAs used were obtained from ThermoFisher (4390843, 4390846, s3585, s3586, s12467, s12468).

### Gene expression analysis by RNA-seq

RNA from myotubes grown in 6-well plates in conditions described above was isolated using the RNeasy Micro Kit from Qiagen (#74004). Quality of RNA was assessed by using a Bioanalyzer 2100 and samples were submitted for library preparation and deep sequencing to the Molecular biology core facility at the Dana Farber Cancer Institute. After sequencing, raw reads of fastq files from all samples were mapped to hg38 genome assemblies using ArrayStudio aligner. Raw read count and FPKM were calculated for all the genes, and DESeq2 was applied to calculate differentially expressed genes using general linear model (GLM). Statistical cutoff of absolute fold change (abs(FC) > 4, FDR < 0.001) were applied to identify differentially expressed protein coding genes. (DATA DEPOSITION INFO TBD)

## RESULTS

### Identification of inhibitors of DUX4 expression

To model FSHD *in vitro*, we differentiated FSHD1 patient-derived immortalized myoblasts into skeletal muscle myotubes. We allowed myoblasts to reach >70% confluency and added differentiation medium lacking growth factors (Figure 1A) (Brewer *et al*., 2008; Krom *et al*., 2012; Thorley *et al*., 2016). After one day of differentiation, we detected DUX4 expression by RT-qPCR and its expression increased throughout the course of myogenic fusion and formation of post-mitotic, multinucleated FSHD myotubes (Figure 1B). Because of the stochastic and low expression levels of DUX4 in FSHD cells, we measured DUX4-regulated genes as an amplified readout of the expression and activity of DUX4. These include *ZSCAN4*, *MBD3L2*, *TRIM43*, *LEUTX* and *KHDC1L* which are among the most commonly described DUX4 targets (Geng *et al*., 2011; Tasca *et al*., 2012; Yao *et al*., 2014; Jagannathan *et al*., 2016; Chen *et al*., 2016; Whiddon *et al*., 2017; Wang *et al*., 2018). These genes were downregulated after DUX4 antisense oligonucleotide treatment of FSHD myotubes and were nearly undetectable or completely absent in FSHD myoblasts or wild-type myotubes (Figure 1C). We concluded that these transcripts were solely dependent on DUX4 expression in differentiating myotubes. Although a number of DUX4-dependent transcripts have been previously described, we selected an assay to specifically detect *MBD3L2* for high-throughput screening because it displayed the best signal window of differential expression in our *in vitro* system comparing FSHD to healthy wildtype myotubes (Figure 1D). With this assay, we identified several small molecules that reduced *MBD3L2* expression after 5 days of differentiation and treatment and showed good reproducibility across replicates (Figure 1E). Validating our results, we found several molecules identified previously to reduce DUX4 expression, including BET inhibitors and β-adrenergic agonists exemplified in Figure S1 (Campbell *et al*., 2017; Cruz *et al*., 2018). However, when treating differentiating FSHD myotubes in our assay, we observed a reduction in fusion as indicated by visual inspection and by the reduction of *MYOG* expression with BET inhibitors. Importantly, we identified multiple scaffolds that inhibit p38 α and β and strongly inhibit the expression of *MBD3L2* without affecting differentiation.

**Figure 1.**
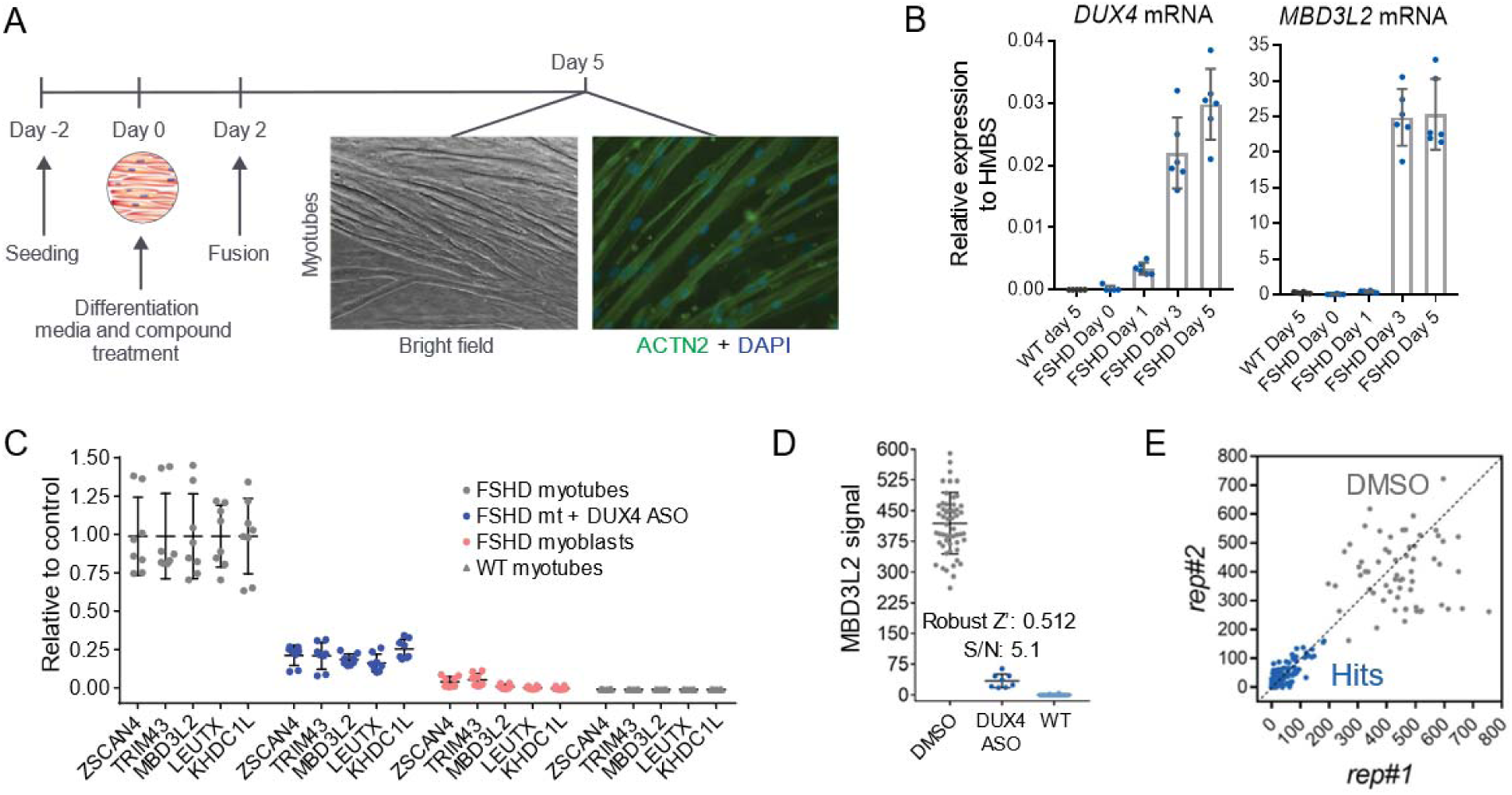
Description of an assay for the identification of inhibitors of DUX4 expression. (**A**) Schematic describing the cellular assay used to identify small molecules that result in the inhibition of DUX4 expression and activity. In short, immortalized FSHD myoblasts (C6, 6.5 D4Z4 RUs) were seeded in 96-well plates 2 days before differentiation was induced. After myoblasts reached confluence, media was replaced and compounds for treatment were added. At day 2, fusion was observed and at day 5, differentiated myotubes were harvested for gene expression analysis or fixed for immunostaining. Representative image of the alpha-actinin staining in differentiated myotubes. (**B**) DUX4 expression is rapidly induced after differentiation of immortalized FSHD myotubes *in vitro*. To measure DUX4 transcript, C6 FSHD myotubes were grown in 12-well plates similarly to A, cells were harvest on day 5 for RNA extraction. RT-qPCR was used to determine expression of *DUX4* mRNA and its downstream gene *MBD3L2* (normalized using *HMBS* as housekeeping). These transcripts were not detected in wild-type immortalized myotubes derived from healthy volunteers. (**C**) Canonical DUX4 target genes are specifically detected in FSHD myotubes and are downregulated when *DUX4* is knocked down using a specific antisense oligonucleotide (ASO). RT-qPCR analysis was used to detect expression in immortalized myoblasts/myotubes. ASO knockdown in FSHD myotubes (mt) was carried out during the 5 days of differentiation. Bars indicate mean±SD. (**D**) A 96-well plate cell-based assay was optimized to screen for inhibitors of DUX4 expression. An assay measuring *MBD3L2* by RT-qPCR was selected because of robust separation and specificity reporting DUX4 activity. *MBD3L2* signal was normalized using *POLR2A* as a housekeeping gene. Bars indicate mean±SD. (**E)** Hits identified in small molecule screen potently reduced the activity of DUX4. X and Y axis show the normalized *MBD3L2* signal obtained from the two replicate wells analyzed.

### p38**α** signaling participates in the activation of DUX4 expression in FSHD myotubes

Potent and selective inhibitors of p38α/β have been previously explored in multiple clinical studies for indications associated with the role of p38α in the regulation of the expression of inflammatory cytokines and cancer (Coulthard *et al*., 2009). We tested several p38α/β inhibitors of different chemical scaffolds in our assays which showed significant inhibition of *MBD3L2* expression (Figure 2A). Importantly, half maximal inhibitory concentrations (IC_50_) obtained for *MBD3L2* reduction were comparable to reported values by other groups in unrelated cell-based assays that measured p38 α/β inhibition, suggesting the specificity for the assigned target(Underwood *et al*., 2000; Campbell *et al*., 2014; Fehr *et al*., 2015). P38α and β kinases phosphorylate a myriad of substrates, including downstream kinases like MAPKAPK2 (also known as MK2) which phosphorylates effector molecules such as heat shock protein 27 (HSP27), as well as a variety of transcription factors including myogenic transcription factors like MEF2C (Simone *et al*., 2004; Knight *et al*., 2012; Segalés, Perdiguero, *et al*., 2016). To determine p38α/β signaling activity in differentiating myoblasts, we measured the levels of phosphorylation of HSP27. As reported previously, we observed increased p38 signaling rapidly upon addition of differentiation media (Figure S2)(Perdiguero *et al*., 2007). We observed P38α/β inhibitors reduced phosphorylated HSP27 levels with similar IC_50_ values to that of *MBD3L2* (Figure 2B). To further validate our findings, we electroporated FSHD myoblasts with siRNAs against p38 α and γ, the most abundant p38 MAPKs in skeletal muscle. After 3 days of differentiation, transient knockdown of p38α showed robust inhibition of expression of *MBD3L2* in FSHD myotubes (Figure 2C) and no significant effects in fusion were observed (Figure S3). We observed that close to 50% reduction of *MAPK14* (p38α) mRNA was sufficient to inhibit *MBD3L2* expression without impacting myogenesis and this level of reduction may account for the differences on myogenesis observed between this study and those previously reported using p38 mouse knockout myoblasts (Perdiguero *et al*., 2007).

**Figure 2.**
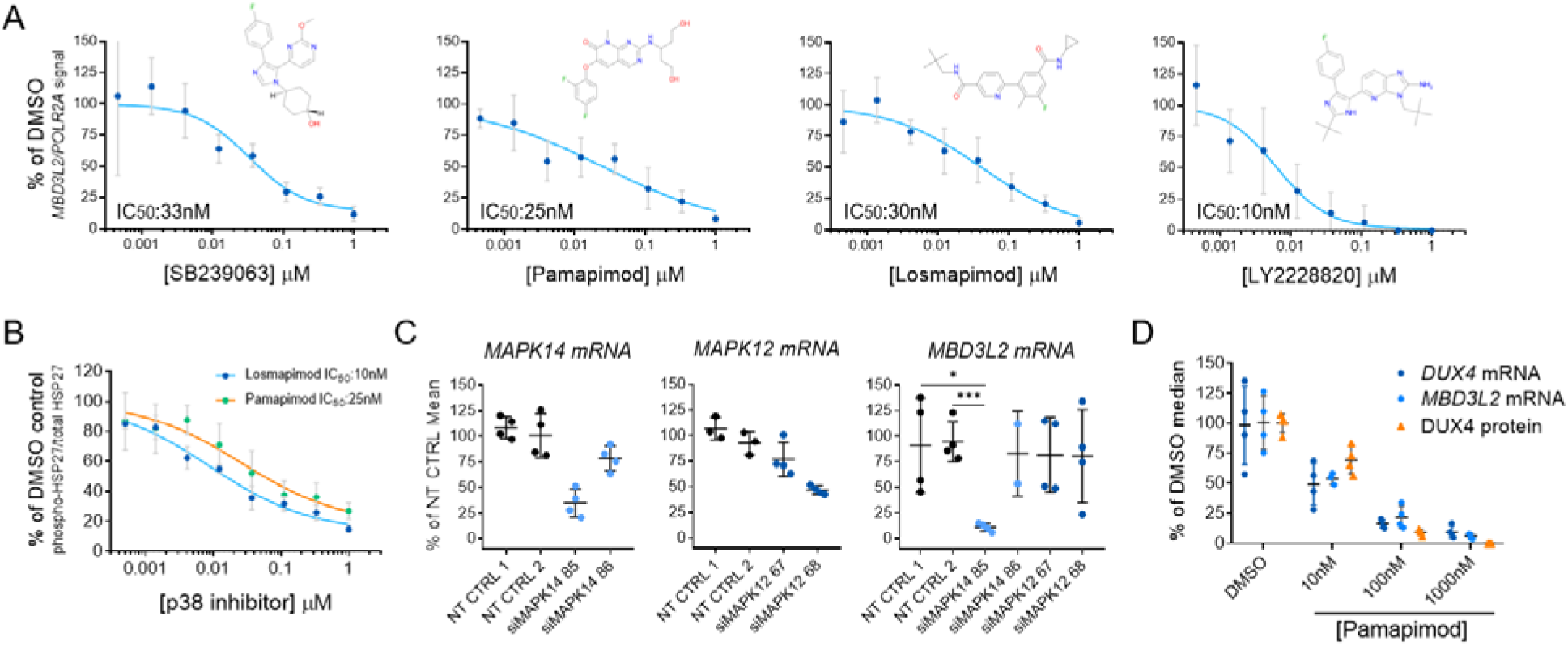
Small molecule inhibitors of p38 alpha reduced expression of DUX4 in FSHD myotubes. (**A**) Diverse inhibitors of p38α/β reduce the expression of *MBD3L2* in differentiating FSHD myotubes. Concentration-dependent responses were observed with all tested inhibitors. Four replicates per concentration were tested to measure reduction of *MBD3L2* in immortalized C6 FSHD myotubes and bars indicate mean±SD. (**B**) P38α/β pathway inhibition in C6 FSHD myotubes. The ratio between phosphorylated HSP27 to total HSP27 was measured by an immunoassay (MSD) after 12h of treatment of C6 FSHD myotubes with the indicated inhibitors. Half maximal inhibitory concentrations (IC_50_) observed for p-HSP27 were comparable to those obtained for reduction of *MBD3L2* expression. Bars indicate mean±SD for four replicate wells. (**C**) Knockdown of p38α (*MAPK14*) results in reduction of *MBD3L2* expression. Immortalized C6 myoblasts were electroporated with siRNAs specific for *MAPK14* (p38α) and *MAPK12* (p38γ) plated and differentiated for 3 days. Expression of the indicated transcripts was measured using RT-qPCR and normalized against *POLR2A*. Reduction of *MBD3L2* expression was observed when >50% knockdown of *MAPK14* was achieved. Bars indicate mean±SD. (**D**) P38α/β inhibition results in the reduction of DUX4 expression. After inhibition, correlated reduction of *DUX4* mRNA, protein and downstream gene *MBD3L2* was observed. To measure DUX4 protein a novel immunoassay was developed using previously described antibodies (see methods and Figure S4). Bars indicate mean±SD, t-test p value * <0.01, *** 0.0002

Our results suggest the p38α pathway is an activator of DUX4 expression in FSHD muscle cells undergoing differentiation. To further understand the reduction in DUX4 expression, we measured the expression of DUX4 transcript and protein upon inhibition of p38 α and β. To measure protein, we developed a highly sensitive assay based on the electrochemiluminescent detection of DUX4 on the Mesoscale Diagnostics (MSD) platform using two previously generated antibodies (Figure S4). We observed that p38α/β inhibition resulted in a highly correlated reduction of DUX4 transcript and protein (Figure 2D). We concluded this led to the reduction in the expression of DUX4 target gene, *MBD3L2*.

### p38 **α** and **β** inhibition normalizes gene expression of FSHD myotubes without impacting the myogenic differentiation program

We further examined the effect of p38 α and β selective inhibition on myotube formation because this pathway has been linked to muscle cell differentiation (Simone *et al*., 2004; Perdiguero *et al*., 2007; Wissing *et al*., 2014; Segalés, Perdiguero, *et al*., 2016; Segalés, Islam, *et al*., 2016). We developed a quantitative assay to measure cell fusion and myotube formation to assess skeletal muscle differentiation *in vitro*. In this assay, we stained immortalized FSHD myotubes cells using antibodies against Myosin Heavy Chains (MHC) and quantified the number of nuclei detected inside MHC-stained region. This provided a way to quantitate the number of cells that successfully underwent the process of *in vitro* myogenesis. P38α/β inhibition by LY2228820 and GW856553X (losmapimod) did not impact differentiation of myoblasts into skeletal muscle myotubes. Treated cells fused properly at all tested drug concentrations to levels comparable to the DMSO control (Figure 3A).

**Figure 3.**
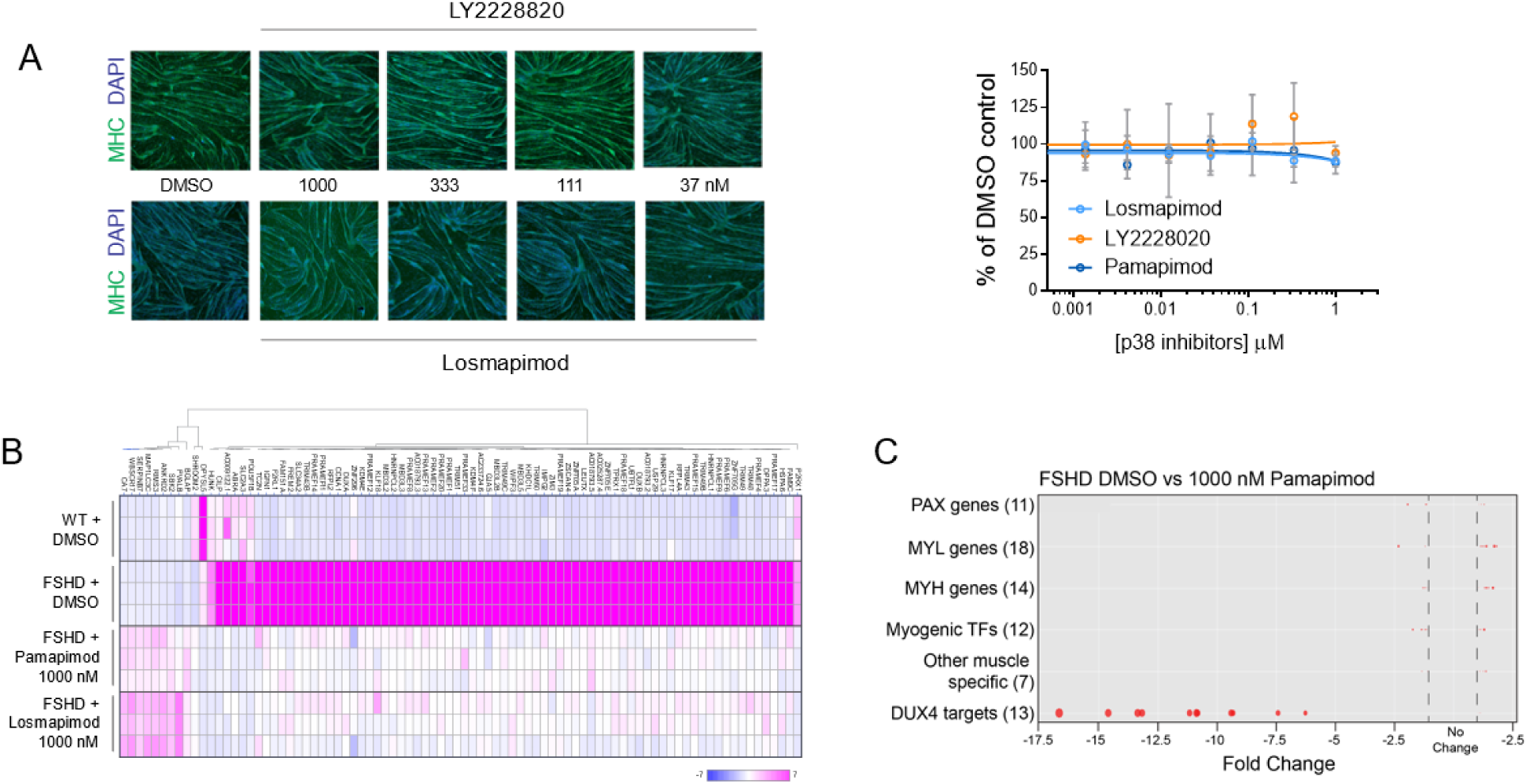
Inhibition of the p38α/β pathway results in normalized gene expression in FSHD myotubes without affecting the differentiation process *in vitro*. (**A**) Quantification of myotube differentiation after p38α/β inhibition. Two inhibitors were used to demonstrate the effects of p38α/β inhibition in a high-content imaging assay to quantify the number of nuclei that properly underwent differentiation by activation of expression of myofiber specific proteins (i.e. MHC). No changes were observed in the morphology of C6 myotubes treated for 5 days. Bars indicate mean±SD. (**B**) Heat map representing fold change of expression levels of differentially expressed genes after p38α/β inhibition in FSHD myotubes for 5 days. 86 genes showed significant changes in expression after treatment with two different inhibitors (abs(FC)>4; FDR<0.001). Each condition was tested in triplicate represented as rows in the heatmap (**C**) DUX4 target genes are specifically downregulated by p38 inhibition. X-axis indicates the fold changes observed in members of the gene families indicated. Diameter of dots represent p-value.

We also further assessed gene expression changes in FSHD myotubes upon p38α/β inhibition. We performed RNA-seq analysis of FSHD and WT myotubes after four days of treatment with vehicle or p38α/β inhibitors. Inhibition of the p38 signaling pathway during differentiation did not induce significant transcriptome changes, and resulted in less than 100 differentially expressed genes (abs(FC)>4; FDR<0.001). About 80% of these differentially expressed genes were known DUX4-regulated transcripts and were all downregulated after p38 α and β inhibition (Figure 3B). This set of DUX4-regulated genes overlapped significantly with genes upregulated in FSHD patient muscle biopsies (Wang *et al*., 2018). Moreover, key driver genes of myogenic programs such as *MYOG*, *MEF* and *PAX* genes and markers of differentiation such as myosin subunits and sarcomere proteins were not affected by p38 inhibition (Figure 3C).

### Inhibition of DUX4 expression results in the reduction of cell death in FSHD myotubes

DUX4 activation and downstream DUX4-regulated target gene expression in muscle cells is toxic, leading to oxidative stress, changes in sarcomere organization, and apoptosis, culminating in reduced contractility, and muscle tissue replacement by fat (Block *et al*., 2013; Bosnakovski *et al*., 2014; Tawil *et al*., 2014; Rickard *et al*., 2015; Homma *et al*., 2015; Choi *et al*., 2016). In particular, apoptotic cells have been detected in skeletal muscle of FSHD patients supporting the hypothesis that programmed cell death is caused by aberrant DUX4 expression and contributes to FSHD pathology (Sandri *et al*., 2001; Statland *et al*., 2015). To test this hypothesis *in vitro*, we evaluated the effect of p38α/β inhibition on apoptosis in FSHD myotubes.

We used an antibody recognizing caspase-3 cleavage products by immunofluorescence to quantify changes in the activation of programmed cell death. Cleavage of caspase-3 is a major step in the execution of the apoptosis signaling pathway, leading to the final proteolytic steps that result in cell death (Fuentes-Prior and Salvesen, 2004; Dix *et al*., 2008; Mahrus *et al*., 2008). We detected activated caspase-3 in FSHD but not in wild-type myotubes and observed a stochastic pattern of expression of DUX4 in FSHD as previously reported (Figure 4A) (Snider *et al*., 2010; Jones *et al*., 2012; van den Heuvel *et al*., 2018). Levels of cleaved caspase-3 were reduced in a concentration-dependent manner with an IC_50_ similar to what we observed for inhibition of the p38 pathway and DUX4 expression (Figure 4B). Moreover, we measured SLC34A2, a DUX4 target gene product using a similar immunofluorescence assay (Figure 3B). This protein was expressed in a similar stochastic pattern observed for active caspase-3 and its expression was also reduced by p38α/β inhibition (Figure 4B and C). Our results demonstrate that DUX4 inhibition in FSHD myotubes results in a significant reduction of apoptosis.

**Figure 4.**
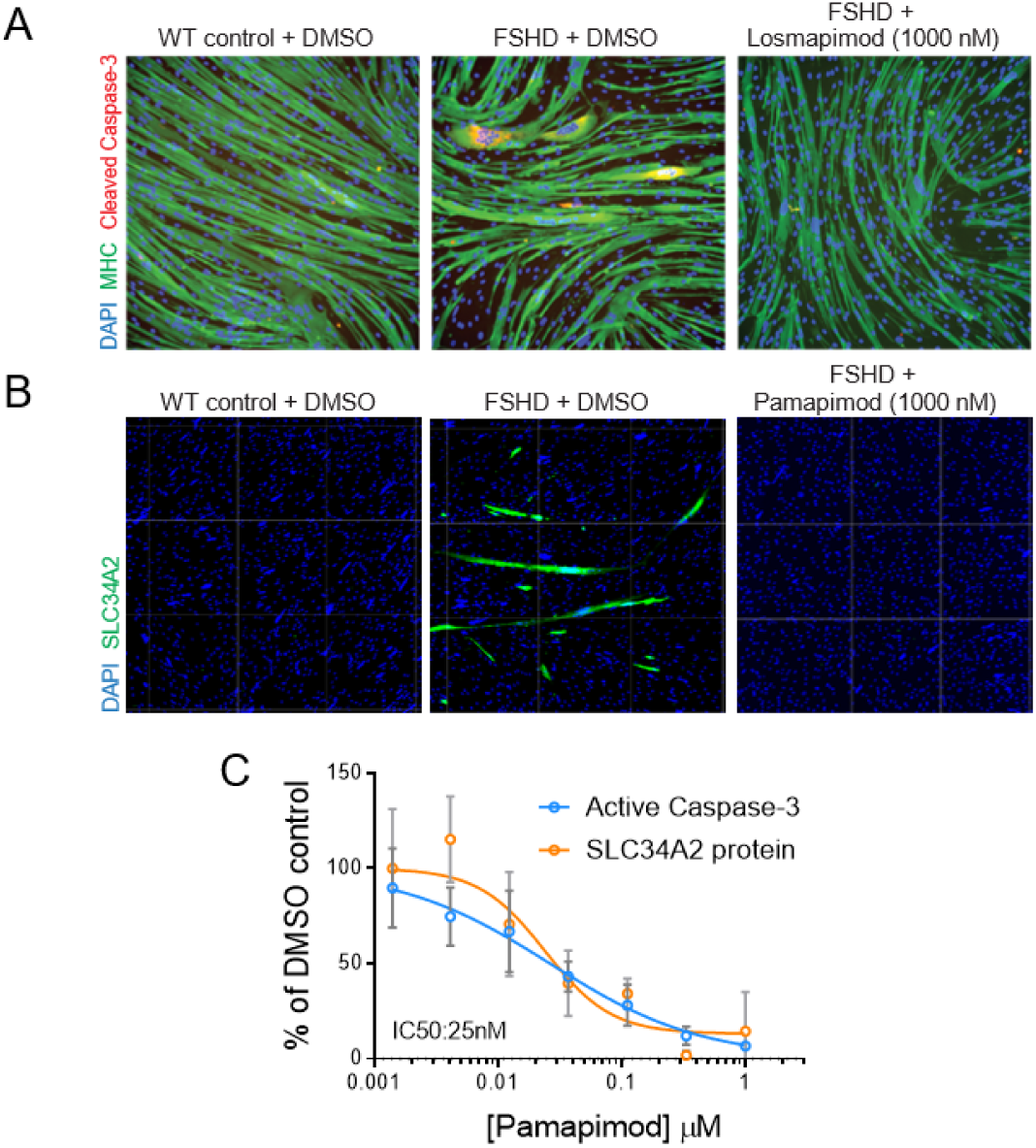
Inhibition of the p38α/β pathway reduced the activation of programmed cell death in differentiating FSHD myotubes. (**A**) A high-content imaging assay was developed to measure cleaved caspase-3 in differentiating myotubes. C6 FSHD myotubes were differentiated and treated for 5 days as indicated above and stained to measure MHC, cleaved-caspase-3 and nuclei. Representative images show that cleaved caspase-3 was only detected in FSHD myotubes, not in wild-type controls or after inhibition of the p38 pathway. Six replicates were imaged and cleaved caspase-3 signal under MHC staining was quantified. (**B**) Stochastic expression of DUX4 target gene, *SLC34A2*, in C6 FSHD myotubes. Expression of SLC34A2 was measured by immunostaining in similar conditions as image above. No expression was detected in wild-type control or p38 inhibitor-treated myotubes. Signal of SLC34A2 under MHC staining was quantified in two replicates (**C**) Concentration-dependent inhibition of the expression of DUX4 target genes is highly correlated to the inhibition of programmed cell death in C6 myotubes. Bars indicate mean±SD.

### p38 **α** and **β** inhibition results in downregulation of DUX4 expression and suppression of cell death across multiple FSHD1 and FSHD2 genotypes

FSHD is caused by the loss of repression at the *D4Z4* locus leading to DUX4 expression in skeletal muscle due to the contraction in the *D4Z4* repeat arrays in chromosome 4 or by mutations in *SMCHD1* and other modifiers such as *DNMT3B*. Primary FSHD myotubes were used to study the *in vitro* efficacy of p38α/β inhibitors across different genotypes. We tested eight FSHD1 primary myoblasts with 2-7 *D4Z4* repeat units and three FSHD2 cell lines with characterized *SMCHD1* mutations. Upon differentiation, the primary cells tested expressed a wide range of *MBD3L2* levels (Figure 5A, number of D4Z4 repeat units or SMCHD1 mutation indicated in parenthesis), comparable to what we and others have observed in other FSHD myotubes (Jones et al., 2012). However, we observed significant inhibition of the DUX4 program expression following treatment with multiple p38α/β inhibitors in all primary myotubes tested from FSHD1 and FSHD2 patients (Figure 5B). Furthermore, this reduction in the DUX4 program resulted in concomitant reduction of cleaved caspase-3 (Figure 5C) without any measurable effects on myotube differentiation (Figure 5D). Our results suggest that the p38α/β pathway critically regulates the activation of DUX4 independently of the mutation driving its expression in FSHD muscle cells.

**Figure 5.**
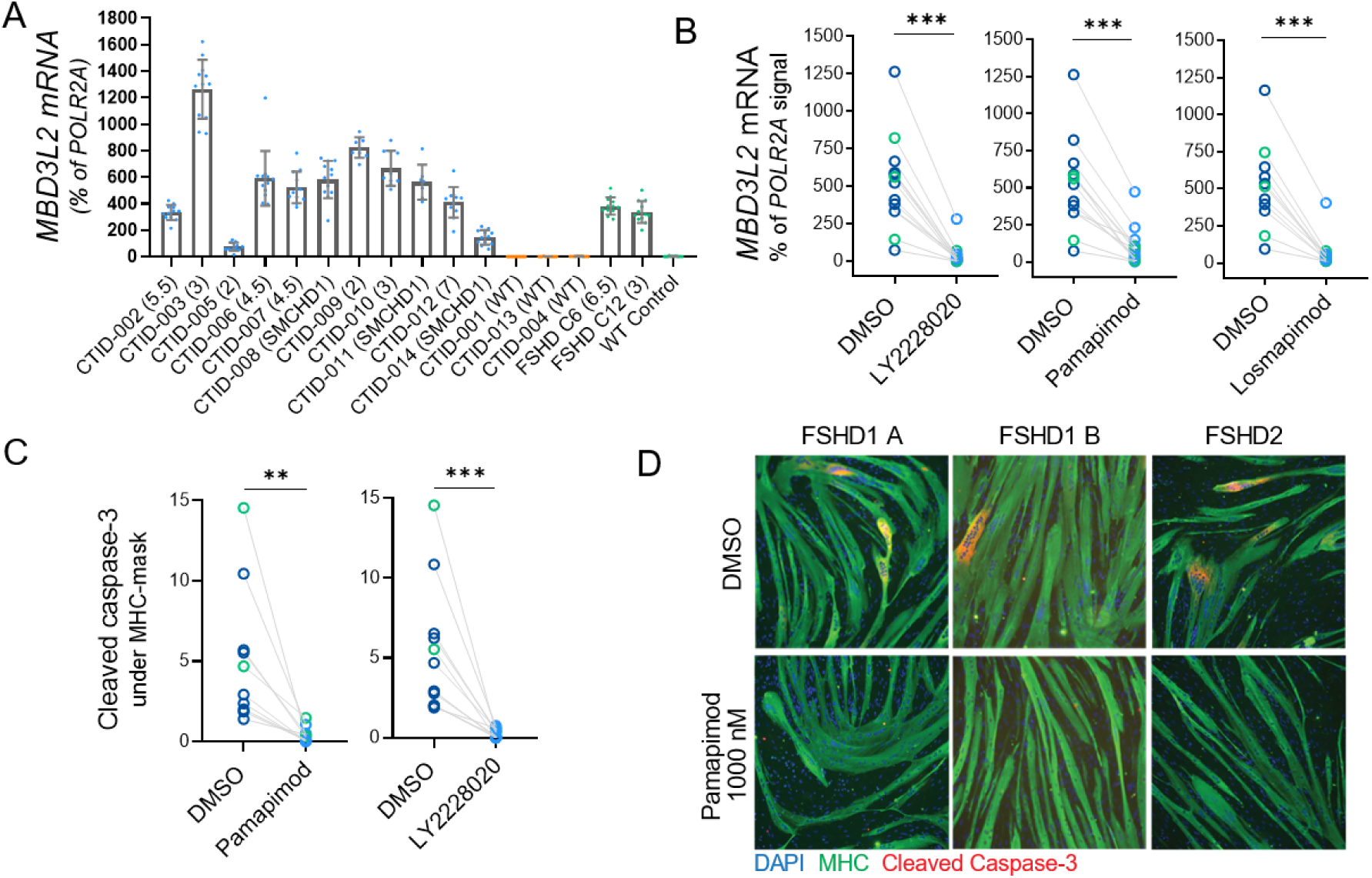
p38α/β inhibition results in the reduction of DUX4 activity and cell death across a variety of genotypes of FSHD1 and FSHD2 primary myotubes. (**A**) Levels of *MBD3L2* expression across different primary and immortalized myotubes determined RT-qPCR. DUX4 activity is only detected in FSHD1/2 lines after 4 days of differentiation. Bars indicate mean±SD and repeat number is indicated in parenthesis in FSHD1 lines and SMCHD1 mutation for FSHD2 lines used. (**B**) Inhibition of the p38α/β pathway results in potent reduction of *MBD3L2* expression activation across the entire set of FSHD primary cells tested. Three different inhibitors were used, and each circle indicates a different FSHD cell line tested. FSHD1 in blue and FSHD2 in green. Expression levels were measured by RT-qPCR in six replicates. (**C and D**) p38α/β pathway inhibition reduces activation of programmed cell death across primary FSHD cell lines with different genotypes. Stochastic activation of caspase-3 in a small number of FSHD myotubes was detected by immunostaining and quantified in all lines. Six replicates were used to quantify signal of cleaved caspase-3 under MHC stained myotubes. Wilconox test, P value **0.002, ***0.0002.

## DISCUSSION

Recent studies have advanced our understanding of the mechanisms that normally lead to the establishment and maintenance of repressive chromatin at the *D4Z4* repeats. Similar to other repetitive elements in somatic cells, chromatin at this locus is decorated by DNA methylation and other histone modifications associated with gene silencing, such as H3K27me3 and H3K9me3 (van Overveld *et al*., 2003; Zeng *et al*., 2009; Cabianca *et al*., 2011; Huichalaf *et al*., 2014; van den Boogaard *et al*., 2016). Factors involved in the deposition of these modifications like *SMCHD1* and *DNMT3B* have been identified by genetic analysis of affected FSHD populations (Lemmers *et al*., 2012; Calandra *et al*., 2016; van den Boogaard *et al*., 2016)ther factors like NuRD and CAF1 have been identified by biochemical approaches isolating proteins that associate with the *D4Z4* locus (Campbell *et al*., 2018). However, sequence-specific transcriptional activators of *DUX4* have remained elusive not only in skeletal muscle but also in the regulation of *DUX4* in the developing embryo, where this factor is normally expressed. Because of the effects of expression of DUX4 in FSHD and the apparent tissue specific expression of DUX4 in skeletal muscle, it has been hypothesized that myogenic regulatory elements upstream of the *D4Z4* repeats regulate the expression of *DUX4* in FSHD (Himeda *et al*., 2014), yet this finding has not led to the identification of other factors that can specifically activate *DUX4*.

In this study, by modelling FSHD *in vitro* and screening a library of probe molecules, we identified p38α as a novel activator of *DUX4* expression in patient-derived FSHD cells. This signaling kinase directly phosphorylates transcription factors involved in myogenesis and may signal directly to activate *DUX4* expression in differentiating myoblasts. Using highly selective and potent small molecules extensively characterized previously, we have studied the pharmacological relationships between the inhibition of this signaling pathway and the inhibition of the expression of DUX4, its downstream gene program expression and its consequences in muscle cells from FSHD patients. These relationships are maintained across multiple FSHD genotypes, including FSHD1 and FSHD2, indicating that this mechanism acts independent of the genetic lesion present in these patients. Our studies show a specific effect of p38 α and β inhibition in downregulation of the DUX4 program and normalization of gene expression compared to cells from healthy donors. Notably, no effects in differentiation were detected at the tested concentrations of p38 inhibitor.

Other recent efforts to identify targets for the treatment of FSHD have reported similar studies in which the investigators followed the expression of *MBD3L2* as a readout for DUX4 expression or by using a reporter driven by the activity of DUX4 in immortalized FSHD myotubes *in vitro* (Campbell *et al*., 2017; Cruz *et al*., 2018). Our results have reproduced their identification of β-adrenergic agonists and BET inhibitors as inhibitors of DUX4 expression. However, these molecules also caused downregulation of the transcription factor *MYOG* expression or affected myoblasts fusion at concentrations similar to the half maximal inhibitory concentration for DUX4 expression inhibition in our model (Figure S1B, lack of fusion indicated by arrow). Recently, an independent study reported that p38α/β inhibitors inhibit expression of DUX4 further validating findings reported here. Importantly in this study, they showed that p38α/β inhibitors are efficacious in downregulating expression of DUX4 in a xenograft mouse model of FSHD.

In humans, previous clinical studies evaluating p38α/β inhibitors in non-FSHD indications under an anti-inflammatory therapeutic hypothesis were tested extensively and shown to be safe and tolerable. However, they never met efficacy endpoints in diseases such as rheumatoid arthritis, chronic obstructive pulmonary disease and acute coronary syndrome (Hill *et al*., 2008; Damjanov *et al*., 2009; Hammaker and Firestein, 2010; Barbour *et al*., 2013; MacNee *et al*., 2013; Norman, 2015; Patnaik *et al*., 2016). Here, we present further evidence from in vitro studies that support the therapeutic hypothesis of treatment of FSHD at its root cause, prevention or reduction of aberrant expression of DUX4, via inhibition of p38 α/β.

## ACKNOWLEDGEMENTS

We thank Peter Jones, Takako Jones and Charis Himeda from the University of Reno for technical advice and guidance during the development of assays in this manuscript and insightful discussions about the regulation of *DUX4* expression. Peter Jones for providing us with constructs for DUX4 overexpression used to validate our DUX4 protein detection assay. Vincent Mouly (Institut Myologie) and Silvère Van der Maarel (LUMC) for providing access to immortalized myoblasts lines. In addition, the authors would like to thank members of Fulcrum Therapeutics for helpful discussions throughout the project. We would also like to thank patients participating in previous studies that have provided tissues to generate cell lines used in this manuscript.

## AUTHORSHIP CONTRIBUTIONS

Participated in research design: Rojas, Valentine, Accorsi, Maglio, Shen, Robertson, Rahl, Kazmirski, Cadavid, Thompson, Tawil, Ronco, Chang, Cacace, Wallace Conducted experiments: Rojas, Valentine, Accorsi, Maglio, Shen, Robertson Contributed new reagents or analytic tools: Valentine, Accorsi, Kazmirski, Tawil Performed data analysis: Rojas, Valentine, Accorsi, Robertson Wrote or contributed to the writing of the manuscript: Rojas, Wallace

## SUPPLEMENTARY FIGURES

**Figure S1.**
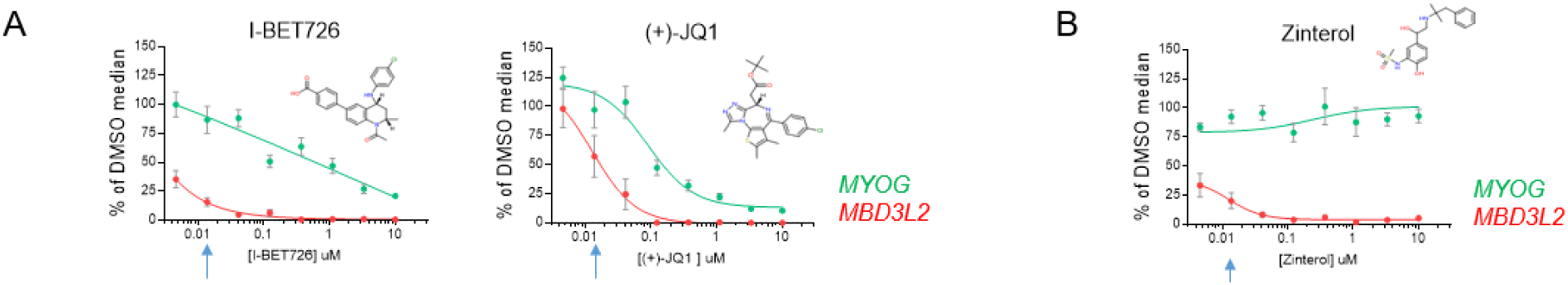
Bromodomain containing proteins inhibitors (A) and β-adrenergic agonist reduced the expression of *MBD3L2* in a concentration dependent manner as previously described (Campbell *et al*., 2017)Arrow indicates concentration at which effects in differentiation started to be observed by visual inspection.

**Figure S2.**
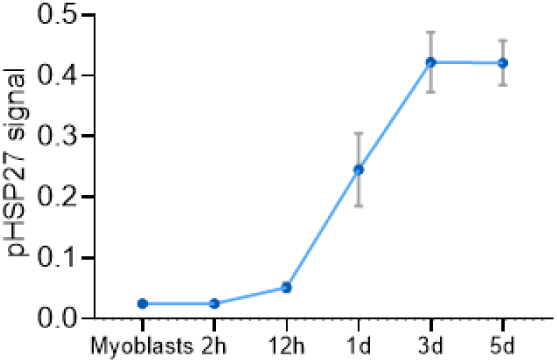
Levels of phosphorylated-HSP27 increase during myogenic differentiation in C6 FSHD myotubes.

**Figure S3.**
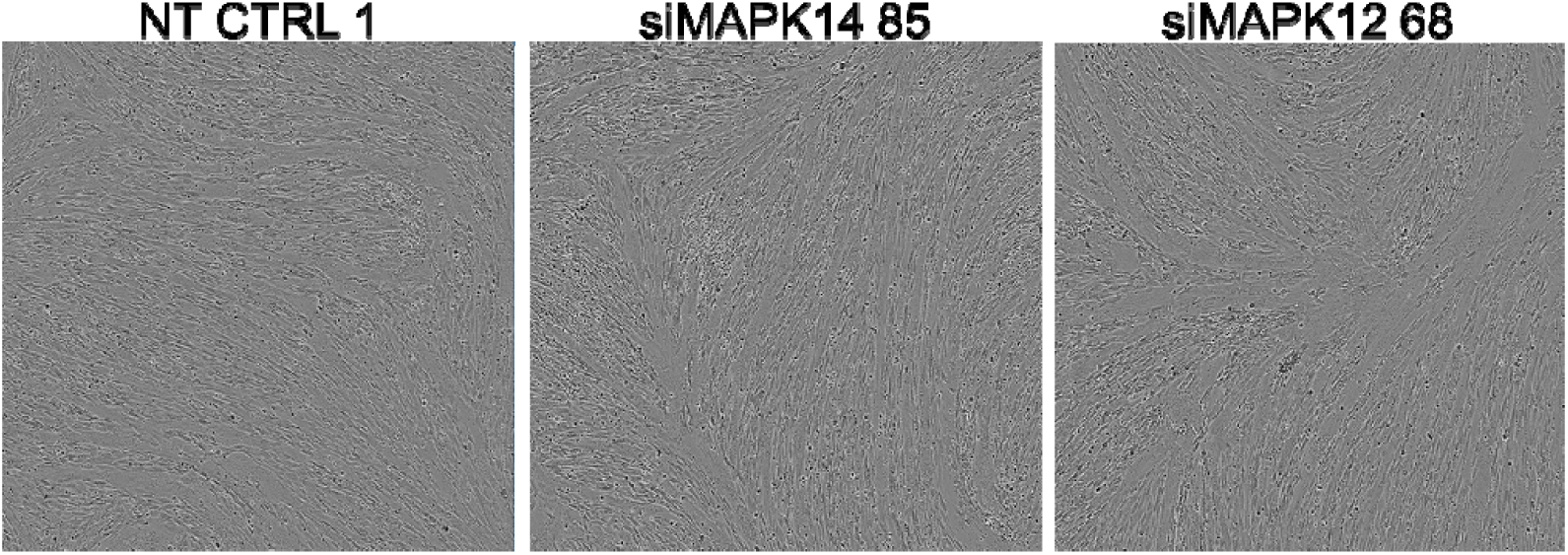
Differentiation of C6 FSHD myotubes was not affected by *MAPK12* and *MAPK14* partial knockdown that resulted in *MBD3L2* level reduction.

**Figure S4.**
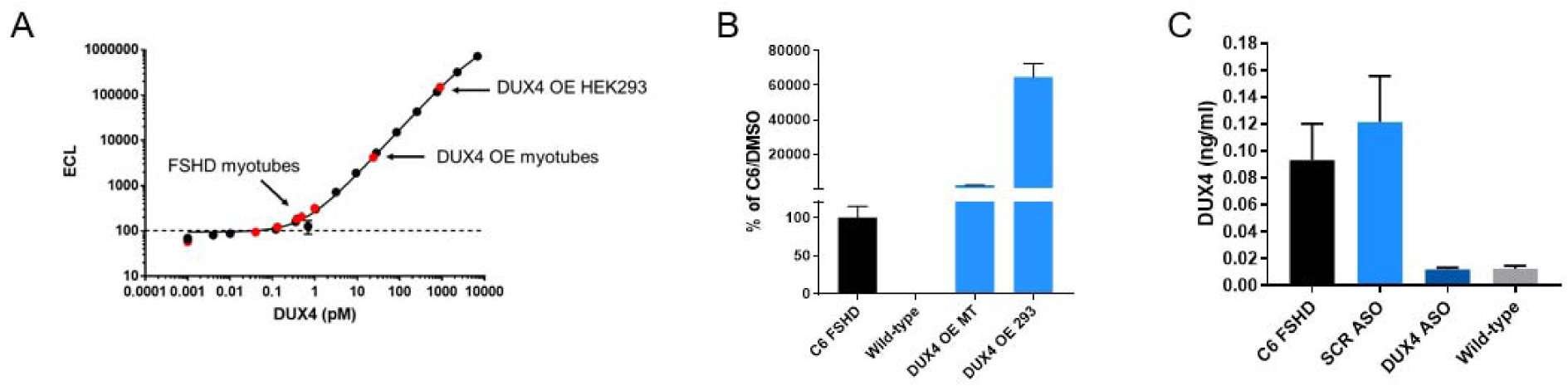
Specific detection of DUX4 protein in mesoscale electro-chemiluminescent immunoassay (A) Recombinant GST-DUX4 calibrator curve. (B) C6 FSHD or wild type 5-day differentiated myotubes, DUX4 overexpressed 1-day differentiated myotubes infected with DUX4 bacmam, DUX4 overexpressed in 293 cells transfected with CMV-DUX4 plasmid. (C) C6 FSHD myotubes treated with scrambled or DUX4 anti-sense oligonucleotide or wild type control.

## REFERENCES

Barbour AM, Saro Blat L, Cai G, Fossler MJ, Sprecher DL, Graggaber J, McGeoch AT, Maison J, and Cheriyan J (2013) Safety, tolerability, pharmacokinetics and pharmacodynamics of losmapimod following a single intravenous or oral dose in healthy volunteers. Brit J Clin Pharmaco 76:99–106.

Block GJ, Narayanan D, Amell AM, Petek LM, Davidson KC, Bird TD, Tawil R, Moon RT, and Miller DG (2013) Wnt/β-catenin signaling suppresses DUX4 expression and prevents apoptosis of FSHD muscle cells. Human Molecular Genetics 22:4661–4672.

Bosnakovski D, Choi S, rasser J, Toso EA, Walters MA, and Kyba M (2014) High-throughput screening identifies inhibitors of DUX4-induced myoblast toxicity. Skeletal Muscle 4:1–11.

Brewer GJ, Boehler MD, Jones TT, and Wheeler BC (2008) NbActiv4 medium improvement to Neurobasal/B27 increases neuron synapse densities and network spike rates on multielectrode arrays. Journal of Neuroscience Methods 170:181–187.

Cabianca DS, Casa V, Bodega B, Xynos A, Ginelli E, Tanaka Y, and Gabellini D (2011) A Long ncRNA Links Copy Number Variation to a Polycomb/Trithorax Epigenetic Switch in FSHD Muscular Dystrophy. Cell 149:819–831.

Calandra P, Cascino I, Lemmers RJ, Galluzzi G, Teveroni E, Monforte M, Tasca G, Ricci E, Moretti F, van der Maarel SM, and Deidda G (2016) Allele-specific DNA hypomethylation characterises FSHD1 and FSHD2. J Med Genet 53:348.

Campbell AE, Oliva J, Yates MP, Zhong J, Shadle SC, Snider L, Singh N, Tai S, Hiramuki Y, Tawil R, van der Maarel SM, Tapscott SJ, and Sverdrup FM (2017) BET bromodomain inhibitors and agonists of the beta-2 adrenergic receptor identified in screens for compounds that inhibit DUX4 expression in FSHD muscle cells. Skeletal Muscle 7:16.

Campbell AE, Shadle SC, Jagannathan S, Lim J-W, Resnick R, Tawil R, van der Maarel SM, and Tapscott SJ (2018) NuRD and CAF-1-mediated silencing of the D4Z4 array is modulated by DUX4-induced MBD3L proteins. Elife 7:e31023.

Campbell RM, Anderson BD, Brooks NA, Brooks HB, Chan EM, Dios A, Gilmour R, Graff JR, Jambrina E, Mader M, McCann D, Na S, Parsons SH, Pratt SE, Shih C, Stancato LF, Starling JJ, Tate C, Velasco JA, Wang Y, and Ye XS (2014) Characterization of LY2228820 Dimesylate, a Potent and Selective Inhibitor of p38 MAPK with Antitumor Activity. Mol Cancer Ther 13:364– 374.

Chen K, Dobson R, Lucet I, Young S, Pearce F, Blewitt M, and Murphy J (2016) The epigenetic regulator Smchd1 contains a functional GHKL-type ATPase domain. Biochemical Journal 473:1733–1744.

Choi SH, Gearhart MD, Cui Z, Bosnakovski D, Kim M, Schennum N, and Kyba M (2016) DUX4 recruits p300/CBP through its C-terminus and induces global H3K27 acetylation changes. Nucleic acids research 44:5161–73.

Coulthard LR, White DE, Jones DL, rmott MF, and Burchill SA (2009) p38MAPK: stress responses from molecular mechanisms to therapeutics. Trends Mol Med 15:369–379.

Cruz JM, Hupper N, Wilson LS, Concannon JB, Wang Y, Oberhauser B, Patora-Komisarska K, Zhang Y, Glass DJ, Trendelenburg A-U, and Clarke BA (2018) Protein kinase A activation inhibits DUX4 gene expression in myotubes from patients with facioscapulohumeral muscular dystrophy. J Biol Chem 293:11837–11849.

Cuenda A, and Rousseau S (2007) p38 MAP-Kinases pathway regulation, function and role in human diseases. Biochimica Et Biophysica Acta Bba - Mol Cell Res 1773:1358–1375.

Damjanov N, Kauffman RS, and Spencer Green GT (2009) Efficacy, pharmacodynamics, and safety of VX 702, a novel p38 MAPK inhibitor, in rheumatoid arthritis: Results of two randomized, double blind, placebo controlled clinical studies. Arthritis & Rheumatism 60:1232–1241.

Deenen JC, Arnts H, van der Maarel SM, Padberg GW, Verschuuren JJ, Bakker E, Weinreich SS, Verbeek AL, and van Engelen BG (2014) Population-based incidence and prevalence of facioscapulohumeral dystrophy. Neurology 83:1056–9.

de Greef JC, Lemmers R, van Engelen B, Sacconi S, Venance SL, Frants RR, Tawil R, and van der Maarel SM (2009) Common epigenetic changes of D4Z4 in contraction dependent and contraction independent FSHD. Hum Mutat 30:1449–1459.

Dion C, Roche S, Laberthonnière C, Broucqsault N, Mariot V, Xue S, Gurzau AD, Nowak A, Gordon CT, Gaillard M-C, El-Yazidi C, Thomas M, Schlupp-Robaglia A, Missirian C, Malan V, Ratbi L, Sefiani A, Wollnik B, Binetruy B, Salort Campana E, Attarian S, Bernard R, Nguyen K, Amiel J, Dumonceaux J, Murphy JM, Déjardin J, Blewitt ME, Reversade B, Robin JD, and Magdinier F (2019) SMCHD1 is involved in de novo methylation of the DUX4-encoding D4Z4 macrosatellite. Nucleic Acids Res 47:gkz005-.

Dix MM, Simon GM, and Cravatt BF (2008) Global Mapping of the Topography and Magnitude of Proteolytic Events in Apoptosis. Cell 134:679–691.

Fehr S, Unger A, Schaeffeler E, Herrmann S, Laufer S, hwab, and Albrecht W (2015) Impact of p38 MAP Kinase Inhibitors on LPS-Induced Release of TNF-α in Whole Blood and Primary Cells from Different Species. Cellular Physiology and Biochemistry 36:2237–2249.

Fuentes-Prior P, and Salvesen GS (2004) The protein structures that shape caspase activity, specificity, activation and inhibition. Biochem J 384:201–232.

Geng LN, Yao Z, Snider L, Fong AP, Cech JN, Young JM, van der Maarel SM, Ruzzo WL, Gentleman RC, Tawil R, and Tapscott SJ (2011) DUX4 Activates Germline Genes, Retroelements, and Immune Mediators: Implications for Facioscapulohumeral Dystrophy. Developmental Cell 22:38–51.

Gordon CT, Xue S, Yigit G, Filali H, Chen K, Rosin N, Yoshiura K, Oufadem M, Beck TJ, McGowan R, Magee AC, Altmüller J, Dion C, Thiele H, Gurzau AD, Nürnberg P, Meschede D, Mühlbauer W, Okamoto N, Varghese V, Irving R, Sigaudy S, Williams D, Ahmed FS, Bonnard C, Kong M, Ratbi I, Fejjal N, Fikri M, Elalaoui S, Reigstad H, Bole-Feysot C, Nitschké P, Ragge N, Lévy N, Tunçbilek G, Teo A, Cunningham ML, Sefiani A, Kayserili H, Murphy JM, Chatdokmaiprai C, Hillmer AM, Wattanasirichaigoon D, Lyonnet S, Magdinier F, Javed A, Blewitt ME, Amiel J, Wollnik B, and Reversade B (2017) De novo mutations in SMCHD1 cause Bosma arhinia microphthalmia syndrome and abrogate nasal development. Nature Genetics, doi: 10.1038/ng.3765.

Hammaker D, and Firestein G (2010) “Go upstream, young man”: lessons learned from the p38 saga. Annals of the Rheumatic Diseases 69:i77–i82.

Hill RJ, Dabbagh K, Phippard D, Li C, Suttmann RT, Welch M, Papp E, Song KW, Chang K, Leaffer D, Kim Y-N, Roberts RT, Zabka TS, Aud D, Porto J, Manning AM, Peng SL, Goldstein DM, and Wong BR (2008) Pamapimod, a Novel p38 Mitogen-Activated Protein Kinase Inhibitor: Preclinical Analysis of Efficacy and Selectivity. Journal of Pharmacology and Experimental Therapeutics 327:610–619.

Himeda CL, Debarnot C, Homma S, Beermann M, Miller JB, Jones PL, and Jones TI (2014) Myogenic Enhancers Regulate Expression of the Facioscapulohumeral Muscular Dystrophy-Associated DUX4 Gene. Mol Cell Biol 34:1942–1955.

Himeda CL, Jones TI, and Jones PL (2015) Facioscapulohumeral Muscular Dystrophy As a Model for Epigenetic Regulation and Disease. Antioxid Redox Sign 22:1463–1482.

Homma S, Beermann M, Boyce FM, and Miller J (2015) Expression of FSHD related DUX4 FL alters proteostasis and induces TDP 43 aggregation. Ann Clin Transl Neurology 2:151–166.

Huichalaf C, Micheloni S, Ferri G, Caccia R, and Gabellini D (2014) DNA Methylation Analysis of the Macrosatellite Repeat Associated with FSHD Muscular Dystrophy at Single Nucleotide Level. PLoS ONE 9:e115278.

Jagannathan S, Shadle S, Resnick R, Snider L, Tawil RN, van der Maarel SM, Bradley RK, and Tapscott SJ (2016) Model systems of DUX4 expression recapitulate the transcriptional profile of FSHD cells. Human molecular genetics, doi: 10.1093/hmg/ddw271.

Jansz N, Chen K, Murphy JM, and Blewitt ME (2017) The Epigenetic Regulator SMCHD1 in Development and Disease. Trends Genet 33:233–243.

Jones T, Chen JC, Rahimov F, Homma S, Arashiro P, Beermann M, King OD, Miller JB, Kunkel LM, Emerson CP, Wagner KR, and Jones PL (2012) Facioscapulohumeral muscular dystrophy family studies of DUX4 expression: evidence for disease modifiers and a quantitative model of pathogenesis. Human Molecular Genetics 21:4419–4430.

Jones TI, King OD, Himeda CL, Homma S, Chen JC, Beermann M, Yan C, Emerson CP, Miller JB, Wagner KR, and Jones PL (2015) Individual epigenetic status of the pathogenic D4Z4 macrosatellite correlates with disease in facioscapulohumeral muscular dystrophy. Clin Epigenetics 7:37.

Jones TI, Yan C, Sapp PC, McKenna-Yasek D, Kang PB, Quinn C, Salameh JS, King OD, and Jones PL (2014) Identifying diagnostic DNA methylation profiles for facioscapulohumeral muscular dystrophy in blood and saliva using bisulfite sequencing. Clin Epigenetics 6:23.

Keren A, Tamir Y, and Bengal E (2006) The p38 MAPK signaling pathway: A major regulator of skeletal muscle development. Mol Cell Endocrinol 252:224–230.

Knight JD, Tian R, Lee RE, Wang F, Beauvais A, Zou H, Megeney LA, Gingras A-C, Pawson T, Figeys D, and Kothary R (2012) A novel whole-cell lysate kinase assay identifies substrates of the p38 MAPK in differentiating myoblasts. Skeletal Muscle 2:1–12.

Krementsov DN, Thornton TM, Teuscher C, and Rincon M (2013) The Emerging Role of p38 Mitogen-Activated Protein Kinase in Multiple Sclerosis and Its Models. Mol Cell Biol 33:3728– 3734.

Krom YD, Dumonceaux J, Mamchaoui K, den Hamer B, Mariot V, Negroni E, Geng LN, Martin N, Tawil R, Tapscott SJ, van Engelen B, Mouly V, Butler-Browne GS, and van der Maarel SM (2012) Generation of Isogenic D4Z4 Contracted and Noncontracted Immortal Muscle Cell Clones from a Mosaic Patient A Cellular Model for FSHD. Am J Pathology 181:1387–1401.

Kyriakis J, and Avruch J (2001) Mammalian mitogen-activated protein kinase signal transduction pathways activated by stress and inflammation. Physiol Rev 81:807–69.

Lemmers RJ, Tawil R, Petek LM, Balog J, Block GJ, Santen GW, Amell AM, van der Vliet PJ, Almomani R, Straasheijm KR, Krom YD, Klooster R, Sun Y, den Dunnen JT, Helmer Q, nlin-Smith C, Padberg GW, van Engelen BG, de Greef JC, Aartsma-Rus AM, Frants RR, de Visser M, Desnuelle C, Sacconi S, Filippova GN, Bakker B, Bamshad MJ, Tapscott SJ, Miller DG, and van der Maarel SM (2012) Digenic inheritance of an SMCHD1 mutation and an FSHD-permissive D4Z4 allele causes facioscapulohumeral muscular dystrophy type 2. Nat Genet 44:1370–1374.

Lemmers RJ, van der Vliet PJ, Klooster R, Sacconi S, Camaño P, Dauwerse JG, Snider L, Straasheijm KR, van Ommen GJ, Padberg GW, Miller DG, Tapscott SJ, Tawil R, Frants RR, and van der Maarel SM (2010) A unifying genetic model for facioscapulohumeral muscular dystrophy. Science (New York, NY) 329:1650–3.

MacNee W, Allan RJ, Jones I, Salvo M, and Tan LF (2013) Efficacy and safety of the oral p38 inhibitor PH-797804 in chronic obstructive pulmonary disease: a randomised clinical trial. Thorax 68:738–745.

Mahrus S, Trinidad JC, Barkan DT, Sali A, Burlingame AL, and Wells JA (2008) Global Sequencing of Proteolytic Cleavage Sites in Apoptosis by Specific Labeling of Protein N Termini. Cell 134:866–876.

Martin E, Bassi R, and Marber (2015) p38 MAPK in cardioprotection – are we there yet? Brit J Pharmacol 172:2101–2113.

Mul K, Lemmers R, Kriek M, van der Vliet PJ, van den Boogaard ML, Badrising UA, Graham JM, Lin AE, Brand H, Moore SA, Johnson K, Evangelista T, Töpf A, Straub V, García S, Sacconi S, Tawil R, Tapscott SJ, Voermans NC, van Engelen B, Horlings C, Shaw ND, and van der Maarel SM (2018) FSHD type 2 and Bosma arhinia microphthalmia syndrome. Neurology 91:e562–e570.

Norman P (2015) Investigational p38 inhibitors for the treatment of chronic obstructive pulmonary disease. Expert Opinion on Investigational Drugs 24:383–392.

Oliva J, Galasinski S, Richey A, Campbell AE, Meyers MJ, Modi N, Zhong J, Tawil R, Tapscott SJ, and erdrup F (2019) Clinically Advanced p38 Inhibitors Suppress DUX4 Expression in Cellular and Animal Models of Facioscapulohumeral Muscular Dystrophy. J Pharmacol Exp Ther 370:219–230.

Patnaik A, Haluska P, Tolcher AW, Erlichman C, Papadopoulos KP, Lensing JL, Beeram M, Molina JR, Rasco DW, Arcos RR, Kelly CS, Wijayawardana SR, Zhang X, Stancato LF, Bell R, Shi P, Kulanthaivel P, Pitou C, Mulle LB, Farrington DL, Chan EM, and Goetz MP (2016) A First-in-Human Phase I Study of the Oral p38 MAPK Inhibitor, Ralimetinib (LY2228820 Dimesylate), in Patients with Advanced Cancer. Clinical Cancer Research 22:1095–1102.

Perdiguero E, Ruiz Bonilla V, Gresh L, Hui L, Ballestar E, Sousa Victor P, Baeza Raja B, Jardí M, Bosch Comas A, Esteller M, Caelles C, Serrano AL, Wagner EF, and Muñoz Cánoves P (2007) Genetic analysis of p38 MAP kinases in myogenesis: fundamental role of p38α in abrogating myoblast proliferation. The EMBO Journal 26:1245–1256.

Rickard AM, Petek LM, and Miller DG (2015) Endogenous DUX4 expression in FSHD myotubes is sufficient to cause cell death and disrupts RNA splicing and cell migration pathways. Hum Mol Genet 24:5901–5914.

Sandri M, Meslemani EA, Sandri C, Schjerling P, Vissing K, Andersen J, Rossini K, Carraro U, and Angelini C (2001) Caspase 3 Expression Correlates With Skeletal Muscle Apoptosis in Duchenne and Facioscapulo Human Muscular Dystrophy. A Potential Target for Pharmacological Treatment? J Neuropathology Exp Neurology 60:302–312.

Segalés J, Islam AB, Kumar R, Liu Q-C, Sousa-Victor P, Dilworth JF, Ballestar E, Perdiguero E, and Muñoz-Cánoves P (2016) Chromatin-wide and transcriptome profiling integration uncovers p38α MAPK as a global regulator of skeletal muscle differentiation. Skeletal Muscle 6:9.

Segalés J, Perdiguero E, and Muñoz-Cánoves P (2016) Regulation of Muscle Stem Cell Functions: A Focus on the p38 MAPK Signaling Pathway. Frontiers in Cell and Developmental Biology 4:91.

Shadle SC, Zhong J, Campbell AE, Conerly ML, Jagannathan S, Wong C-J, Morello TD, van der Maarel SM, and Tapscott SJ (2017) DUX4-induced dsRNA and MYC mRNA stabilization activate apoptotic pathways in human cell models of facioscapulohumeral dystrophy. PLOS Genetics 13:e1006658.

Shaw ND, Brand H, Kupchinsky ZA, Bengani H, Plummer L, Jones TI, Erdin S, Williamson KA, Rainger J, Stortchevoi A, Samocha K, Currall BB, Dunican DS, Collins RL, Willer JR, Lek A, Lek M, Nassan M, Pereira S, Kammin T, Lucente D, Silva A, abra C, Chiang C, An Y, Ansari M, Rainger JK, Joss S, Smith J, Lippincott MF, Singh SS, Patel N, Jing JW, Law JR, Ferraro N, Verloes A, Rauch A, Steindl K, Zweier M, Scheer I, Sato D, Okamoto N, Jacobsen C, Tryggestad J, Chernausek S, Schimmenti LA, Brasseur B, Cesaretti C, García-Ortiz JE, Buitrago T, Silva O, Hoffman JD, Mühlbauer W, Ruprecht KW, Loeys BL, Shino M, Kaindl AM, Cho C-H, Morton CC, Meehan RR, van Heyningen V, Liao EC, Balasubramanian R, Hall JE, Seminara SB, Macarthur D, Moore SA, Yoshiura K, Gusella JF, Marsh JA, Jr JM, Lin AE, Katsanis N, Jones PL, Jr WF, Davis EE, FitzPatrick DR, and Talkowski ME (2017) SMCHD1 mutations associated with a rare muscular dystrophy can also cause isolated arhinia and Bosma arhinia microphthalmia syndrome. Nature Genetics, doi: 10.1038/ng.3743.

Simone C, Forcales S, Hill DA, Imbalzano AN, Latella L, and Puri P (2004) p38 pathway targets SWI-SNF chromatin-remodeling complex to muscle-specific loci. Nat Genet 36:738–743.

Snider L, Geng LN, Lemmers R, Kyba M, Ware CB, Nelson AM, Tawil R, Filippova GN, van der Maarel SM, Tapscott SJ, and Miller DG (2010) Facioscapulohumeral Dystrophy: Incomplete Suppression of a Retrotransposed Gene. PLoS Genetics 6:e1001181.

Statland JM, Odrzywolski KJ, Shah B, Henderson D, Fricke AF, van der Maarel SM, Tapscott SJ, and Tawil R (2015) Immunohistochemical Characterization of Facioscapulohumeral Muscular Dystrophy Muscle Biopsies. J Neuromuscul Dis 2:291–299.

Statland JM, and Tawil R (2014) Risk of functional impairment in Facioscapulohumeral muscular dystrophy. Muscle Nerve 49:520–527.

Tasca G, Pescatori M, Monforte M, Mirabella M, Iannaccone E, Frusciante R, Cubeddu T, Laschena F, Ottaviani P, and Ricci E (2012) Different Molecular Signatures in Magnetic Resonance Imaging-Staged Facioscapulohumeral Muscular Dystrophy Muscles. PLoS ONE 7:e38779.

Tawil R, Forrester J, Griggs RC, Mendell J, Kissel J, rmott M, King W, Weiffenbach B, and Figlewicz D (1996) Evidence for anticipation and association of deletion size with severity in facioscapulohumerd muscular dystrophy. Ann Neurol 39:744–748.

Tawil R, Kissel JT, Heatwole C, Pandya S, Gronseth G, and Benatar M (2015) Evidence-based guideline summary. Neurology 85:357–364.

Tawil R, van der Maarel SM, and Tapscott SJ (2014) Facioscapulohumeral dystrophy: the path to consensus on pathophysiology. Skeletal Muscle 4:1–15.

Thorley M, Duguez S, Mazza E, Valsoni S, Bigot A, Mamchaoui K, Harmon B, Voit T, Mouly V, and Duddy W (2016) Skeletal muscle characteristics are preserved in hTERT/cdk4 human myogenic cell lines. Skeletal Muscle 6:43.

Underwood D, Osborn R, Kotzer C, Adams J, Lee J, Webb E, Carpenter D, Bochnowicz S, Thomas H, Hay D, and Griswold D (2000) SB 239063, a potent p38 MAP kinase inhibitor, reduces inflammatory cytokine production, airways eosinophil infiltration, and persistence. J Pharmacol Exp Ther 293:281–8.

van den Boogaard ML, Lemmers R, Balog J, Wohlgemuth M, Auranen M, Mitsuhashi S, van der Vliet PJ, Straasheijm KR, van den Akker R, Kriek M, Laurense-Bik M, Raz V, van Ostaijen-ten Dam MM, Hansson K, van der Kooi EL, Kiuru-Enari S, Udd B, van Tol M, Nishino I, Tawil R, Tapscott SJ, van Engelen B, and van der Maarel SM (2016) Mutations in DNMT3B Modify Epigenetic Repression of the D4Z4 Repeat and the Penetrance of Facioscapulohumeral Dystrophy. Am J Hum Genetics 98:1020–1029.

van den Boogaard ML, Lemmers R, Camaño P, van der Vliet PJ, Voermans N, van Engelen BG, de Munain A, Tapscott SJ, van der Stoep N, Tawil R, and van der Maarel SM (2015) Double SMCHD1 variants in FSHD2: the synergistic effect of two SMCHD1 variants on D4Z4 hypomethylation and disease penetrance in FSHD2. Eur J Hum Genet 24:78–85.

van den Heuvel A, Mahfouz A, Kloet SL, Balog J, van Engelen BG, Tawil R, Tapscott SJ, and van der Maarel SM (2018) Single-cell RNA-sequencing in facioscapulohumeral muscular dystrophy disease etiology and development. Hum Mol Genet, doi: 10.1093/hmg/ddy400.

van Overveld PG, Lemmers RJ, Sandkuijl LA, Enthoven L, Winokur ST, Bakels F, Padberg GW, van Ommen G-JB, Frants RR, and van der Maarel SM (2003) Hypomethylation of D4Z4 in 4q-linked and non-4q-linked facioscapulohumeral muscular dystrophy. Nat Genet 35:ng1262.

Viemann D, Goebeler M, Schmid S, Klimmek K, Sorg C, Ludwig S, and Roth J (2004) Transcriptional profiling of IKK2/NF-κB— and p38 MAP kinasedependent gene expression in TNF-α—stimulated primary human endothelial cells. Blood 103:3365–3373.

Wang LH, Friedman SD, Shaw D, Snider L, Wong C-J, Budech CB, Poliachik SL, Gove NE, Lewis LM, Campbell AE, Lemmers RJ, Maarel SM, Tapscott SJ, and Tawil RN (2018) MRI-informed muscle biopsies correlate MRI with pathology and DUX4 target gene expression in FSHD. Hum Mol Genet, doi: 10.1093/hmg/ddy364.

Whiddon JL, Langford AT, Wong C-J, Zhong J, and Tapscott SJ (2017) Conservation and innovation in the DUX4-family gene network. Nature Genetics, doi: 10.1038/ng.3846.

Whitmarsh AJ (2010) A central role for p38 MAPK in the early transcriptional response to stress. Bmc Biol 8:1–3.

Wissing ER, Boyer JG, Kwong JQ, Sargent MA, Karch J, McNally EM, Otsu K, and Molkentin JD (2014) P38α MAPK underlies muscular dystrophy and myofiber death through a Bax-dependent mechanism. Human Molecular Genetics 23:5452–5463.

Yao Z, Snider L, Balog J, Lemmers R, Maarel SM, Tawil R, and Tapscott SJ (2014) DUX4-induced gene expression is the major molecular signature in FSHD skeletal muscle. Hum Mol Genet 23:5342–5352.

Zarubin T, and Han J (2005) Activation and signaling of the p38 MAP kinase pathway. Cell Res 15:7290257.

Zeng W, de Greef JC, Chen Y-Y, Chien R, Kong X, Gregson HC, Winokur ST, Pyle A, Robertson KD, Schmiesing JA, Kimonis VE, Balog J, Frants RR, Ball AR, Lock LF, Donovan PJ, van der Maarel SM, and Yokomori K (2009) Specific Loss of Histone H3 Lysine 9 Trimethylation and HP1γ/Cohesin Binding at D4Z4 Repeats Is Associated with Facioscapulohumeral Dystrophy (FSHD). Plos Genet 5:e1000559.

Zetser A, Gredinger E, and Bengal E (1999) p38 Mitogen-activated Protein Kinase Pathway Promotes Skeletal Muscle Differentiation PARTICIPATION OF THE MEF2C TRANSCRIPTION FACTOR. J Biol Chem 274:5193–5200.

